# Characterisation of behaviours relevant to apathy syndrome in the aged male rat

**DOI:** 10.1101/2024.02.02.578556

**Authors:** Megan G Jackson, Stafford L Lightman, Emma S J Robinson

## Abstract

Apathy is a complex psychiatric syndrome characterised by motivational deficit, emotional blunting and cognitive changes. It occurs alongside a broad range of neurological disorders, but also occurs in otherwise healthy ageing. Despite its clinical prevalence, apathy does not yet have a designated treatment strategy. Generation of a translational animal model of apathy syndrome would facilitate the development of novel treatments. Given the multidimensional nature of apathy, a model cannot be achieved with a single behavioural test. Using a battery of behavioural tests we investigated whether aged rats exhibit behavioural deficits across different domains relevant to apathy.

Using the effort for reward and progressive ratio tasks we found that aged rats (21-27 months) show intact reward motivation. Using the novelty supressed feeding test and position-based object exploration we found aged rats showed increased anxiety-like behaviour inconsistent with emotional blunting. The sucrose preference test and reward learning assay showed intact reward sensitivity and reward-related cognition in aged rats. However, using a bowl-digging version of the probabilistic reversal learning task, we found a deficit in cognitive flexibility in aged rats that did not translate across to a touchscreen version of the task.

While these data reveal important changes in cognitive flexibility and anxiety associated with ageing, aged rats do not show deficits across other behavioural domains relevant to apathy. This suggests that aged rats are not a suitable model for age-related apathy syndrome. These findings contrast with previous work in mice, revealing important species differences in behaviours relevant to apathy syndrome in ageing.

## 1. Introduction

Apathy is a complex neuropsychiatric syndrome with symptoms spanning emotional, cognitive and behavioural domains (Levy and Dubois, 2006, Robert et al., 2018). It accompanies a broad range of neurodegenerative, neurocognitive and psychiatric disorders, but also occurs in otherwise healthy ageing (Brodaty et al., 2009). Apathy is associated with higher mortality, poorer quality of life, cognitive decline and significant caregiver stress (Ishii et al., 2009, Gerritsen et al., 2005). It can also negate the positive effects of therapeutic intervention by affecting adherence to medication and healthcare appointments (Padala et al., 2008). Despite its clinical relevance and research importance, the underlying neurobiology of apathy remains unclear, and does not yet have a designated treatment strategy.

Generating a translational animal model for apathy could allow for potential treatments to be tested and mechanisms to be elucidated. However, comprehensive models of apathy syndrome are so far limited and most studies rely on a single behavioural test. Given the multidimensional nature of the syndrome, an apathy model is unlikely to be achieved with a single behavioural test. At the preclinical level, a deficit in motivation for reward alongside evidence of emotional blunting represent behavioural hallmarks of apathy behaviour and distinguish it from behaviours relating to major depressive disorder, which is often characterised by negative affective state and exaggerated response to negative stimuli (Jackson and Robinson, 2022, Beck, 1967). In a behavioural battery, previous work has shown that aged mice show deficits in physical effort-based motivation for reward in the effort for reward (EfR) and progressive ratio (PR) tasks and evidence of emotional blunting (reduction in anxiety response, reduced corticosterone response and brain activation in stress areas following a restraint stressor) (Jackson et al., 2021) suggesting natural ageing could be used as a model of apathy. In addition to physical effort and emotional processing changes, patients may experience cognitive symptoms, including a reduction in cognitive flexibility, planning and decision making (Malpetti et al., 2021). The probabilistic reversal learning task (PRLT) provides a translational measure of cognitive flexibility and feedback sensitivity used across species (Bari et al., 2010, Der-Avakian et al., 2013). In this paradigm, subjects must adjust their behaviour to changing contingencies. This task is sensitive to deficits in cognitive flexibility in conditions where apathy commonly occurs, such as Parkinson’s Disease (Peterson et al., 2009) as such, use of this task may be a relevant measure of cognitive deficits associated with apathy syndrome at the preclinical level.

While we have previously demonstrated that aged mice may be suitable to assess certain behaviours relating to distinct apathy domains, relevant cognitive changes have not been well characterised. Rats may be a more suitable species for tasks of this nature given their greater propensity to reverse in standard probabilistic paradigms and more established repertoire of cognitive tasks (Metha et al., 2020). Therefore, in this study we combined tests of reward motivation (including the effort for reward, progressive ratio task) and emotional processing (novelty supressed feeding test (NSFT), restraint stress, position-based object exploration, reward learning assay (RLA), sucrose preference test (SPT)) in addition to measures of cognitive flexibility (probabilistic reversal learning) to investigate whether aged rats exhibit behavioural deficits across different domains relevant to apathy syndrome. For the purpose of this study we defined aged rats as animals more than 21 months old.

## 2. Methods

### 2.1 Subjects

10 aged, male Sprague-Dawley rats (aged 21 months old at start of experiments, weighing 467-530g and 25 months, weighing 480-555g by end) and 10 sex and strain-matched younger controls (4 months old and weighing 310-320 g at start of experiments, 8 months, weighing 470-545g by end) were kindly supplied by Eli Lilly. Sample size was based on detecting a large effect size (∼ 1.0) with mean difference and variance based on previous behavioural studies using a similar task and using aged animals (Jackson et al., 2021). The aged rats had spent a period of approx. 5 months on a restricted diet (16g fed between ZT2-3) at Eli Lilly. This was continued for a period of 5 days at Bristol to aid adjustment before being put on *ad libitum* feeding for the beginning of experiments. Rats were housed in enriched cages, consisting of a red shelf, wooden chew block and cardboard tube in groups of either threes or pairs (counterbalanced across age groups). Water and food were given *ad libitum*, unless undergoing operant training, novelty supressed feeding testing or sucrose preference. They were kept in 12:12 reverse lighting, with lights OFF at 8.15am and ON at 8.15pm. Behavioural testing was conducted under red lighting and within the animal’s active phase. Due to age being the factor of analysis, rodents could not be fully randomised to groups. However, within a behavioural study, order of testing was counterbalanced across age to mitigate time of day effects. Aged animals were checked daily for general health and any impairments which could confound interpretation of behavioural studies. Any animal which showed signed of declining health was removed from the study. It should be noted that behavioural tests were performed sequentially and therefore there is the potential that advancing age differentially affected some measures (timeline provided in supplementary methods). A summary of the behavioural tasks and associated behavioural domains tested are provided in **table 1**. To help reduce confounds associated with the order of the behavioural tasks, the design randomised the tasks between measures of motivation-related and emotional behaviour. Due to clear visual differences between young and aged rats, the experiment could not be blind to age while running behavioural experiments. All experiments took place in the animals’ active phase and were performed in accordance with the Animals (Scientific Procedures) Act (United Kingdom) 1986 and were approved by the University of Bristol Animal Welfare and Ethical Review Body (AWERB).

**Table 1.** Summary of behavioural tasks used, the associated Research Domain Criteria behavioural construct tested (Cuthbert, 2015, Jackson and Robinson, 2022) and hypothesised mapping to apathy symptom domains according to the 2018 consensus group classification (Robert et al., 2018). Apathy syndrome consists of behavioural changes relating to multiple behavioural domains including ‘emotion’, ‘behaviour-cognition’ and ‘social motivation’. In patients, behaviours relating to the emotion domain include a blunted or lack of reaction to an external piece of news (positively or negatively valanced). Behaviours relating to the behaviour-cognition domain include reduced motivation to start/finish a task. Behaviours relating to the social motivation domain include a reduction or lack of initiation of social contact (Ang et al., 2017).

| Behavioural task (rodent) | Behavioural/physiological construct(s) measured (Research Domain Criteria) | Hypothesised mapping to apathy domain (2018 international consensus group classification) |
| --- | --- | --- |
| Position-based object exploration | Spontaneous exploration, anxiety-like behaviour | Behaviour-cognition<br>Emotion |
| Novelty suppressed feeding test | Anxiety-like behaviour | Emotion |
| Effort for reward | Motivation for reward | Behaviour-cognition |
| Progressive ratio | Motivation for reward | Behaviour-cognition |
| Restraint stress | Hormonal reactivity to stress | Emotion |
| Reward learning assay | Reward learning, affective state | Emotion |
| Sucrose preference test | Reward sensitivity | Emotion |
| Probabilistic reversal learning (bowl digging) | Cognitive flexibility, feedback sensitivity | Behaviour-cognition |
| Probabilistic reversal learning (touchscreen) | Cognitive flexibility, feedback sensitivity | Behaviour-cognition |

To further manage the impacts of health impairment confounding behaviour interpretation, a second cohort of rats were brought in (**supplementary figures, S1**). Here, a further 10 aged male Sprague-Dawley rats were obtained from Eli Lilly (aged 23 months at experiment onset, weighing 440-525g and aged 27 months and weighing 401-560g by experiment end). Rats arrived at the Bristol unit aged 21 months but due to COVID restrictions experiments did not start until 2 months later. During this delay period aged rats were fed a restricted diet of 18g/rat. One week before experimentation started aged rats were put onto *ad libitum* feeding to match young controls. 10 strain and sex-matched controls were obtained from Envigo at a weight between 300-324g, aged approx. 3 months and aged 7 months and weighing between 403-460g by experiment end.

### 2.2 Object exploration

Rats were placed in a square arena (1m x 1m) lined with sawdust. They were allowed to explore the arena freely under red lighting for a period of 15 min. The following day, a novel object (either a pinecone or a scrubbing brush) was placed in either the centre (higher risk) or to the side of the arena (lower risk). The rat was placed in the arena and could explore the arena and novel object for a period of 10 mins. This test is based on the well-characterized principle that a rodent will exhibit thigmotaxis when placed in an open arena (Hall and Ballachey, 1932). As such, the rodent should bias its exploration of an object positioned to the side of the arena versus the object placed in the center. Sessions took place over two days, and position and object were counterbalanced over these two sessions and across age group. Their movement was recorded using a Logitech webcam, and the videos were subsequently coded so that the experimenter was blind to age group. Time spent exploring the object and bouts of exploration were manually recorded, and a % object placement preference was calculated.

### 2.3 Novelty supressed feeding test

Rats were food restricted for 24 hrs before onset of the test. During testing, rats were placed in a circular arena (70 cm in diameter), lined with sawdust. The test was conducted under white lighting, but during the animals’ active phase. A bowl was placed in the centre of arena containing standard lab chow. Time taken for the rat to approach the bowl and to eat from it was manually recorded using a stopwatch. Sawdust was redistributed between rats and the bowl was washed and fresh food pellets were used each time.

To test for potential age-related appetite confounds that may drive results observed in the novelty supressed feeding test, rats were food restricted for 24hrs. They were then placed in individual test cages they had previously been habituated to, with a known weight of food. After a period of 10 mins, total amount eaten in g was measured. Rats were then returned to *ad libitum* feeding.

### 2.4 Lever training and fixed ratio testing

Rats were trained in sound-proofed operant boxes (Med Associates Inc) which were run on Klimbic software (Conclusive Solutions Ltd., UK) similar to the protocol previously described by (Griesius et al., 2020). Each box consisted of two levers positioned either side of a reward magazine which was connected to a reward pellet dispenser (45 mg rodent tablet, TestDiet, Sandown) (see chapter 2, fig 2A). Only one lever was active during training/testing, and positioning was counterbalanced across cohorts. Unless otherwise stated, a session lasted 30 mins. Rats first learned to associate the magazine with the delivery of reward where pellets were dispensed automatically every 40 s for two sessions. Rats then underwent three sessions of continuous reinforcement learning, in which they learned to associate a lever press with the delivery of a reward. They then progressed on to fixed ratio (FR) training. FR refers to the number of lever presses to receive one reward pellet. Rats progressed through FR1, 2, 4 and 8 after two consecutive sessions of 100+ trials, and then FR16 after two consecutive sessions of 70+ trials. All rats then underwent at least 5 additional FR16 sessions.

### 2.5 Effort for reward task

Directly following lever training and testing up to FR16, rats underwent a single session of the EfR task, previously described by (Griesius et al., 2020, Salamone et al., 1991). Here, rats underwent a standard FR16 session, with the addition of a bowl of powdered standard lab chow placed in front of the inactive lever. The chow was accessed through a small hole in the lid. The FR16 requirement for a reward pellet represents the high effort, high value reward option, while the freely available lab chow represents the low effort, low value reward option. Throughout the 30 min session the animal could freely choose to work for reward or consume the chow. At a later timepoint in the experimental timeline, rats completed 5 consecutive EfR sessions under both food restriction and *ad libitum*. The primary output measures for this task were the number of high effort, high reward trials, later termed ‘trials’, and consumption of the low effort, low value reward (standard powdered lab chow).

### 2.6 Progressive ratio task

Rats were tested in the PR task, using a response schedule of F(*n*) = 5 × EXP(0.2*n*)−5 (Roberts and Richardson, 1992, Jackson et al., 2021). The session lasted a maximum of 60 minutes. The response ratio completed before the rat stopped responding within the session time limit for a period of 5 minutes was used as a breakpoint. In some cases, rats continued until the max time limit and this was termed ‘final ratio completed’ and was analysed separately from breakpoint. A separate PR test session was conducted under food restriction conditions and *ad libitum* feeding. The primary output measures for this task were breakpoint and/or final ratio completed.

### 2.7 Restraint stress and corticosterone analysis

Rats were restrained for 30 mins using a restraint tube (52-0292 RODENT RESTRAINER, Biochrom). Blood samples were collected immediately preceding and post-stress. Blood was collected by inserting a butterfly needle into the tail vein. Blood was dripped into an Eppendorf containing 50 µL EDTA. Blood samples were then centrifuged at 8000 rpm for 10 mins. Plasma was collected and stored at -20 °c until analysis. A radioimmunoassay (RIA) was performed in-house to measure total corticosterone from restraint stress blood samples as previously described (Windle et al., 1998). Samples were analysed by RIA in triplicate to account for variation in assay results and the median value was used in analysis. See supplementary methods for more details. The primary output measure was corticosterone (ng/ml) pre- and post-stress.

### 2.8 Sucrose preference test

During habituation, rats were water restricted for a period of 4 h and were then placed in test cages containing two sipper sacks (Avidity Science) for 1 h. On the first and third day the sipper sacks contained 2 % sucrose, and on the second day they contained tap water. On the test day, the rat was presented with one sipper sack containing 2 % sucrose and the other containing tap water. Starting position of sucrose was counter balanced and swapped at 30 mins. Amount consumed (ml) was measured at each time point and a % sucrose preference was calculated as primary output measure using: (amount of sucrose solution consumed/total solution consumed) x 100.

### 2.9 Bowl digging training

Rats were first habituated to a 40 cm x 40 cm Perspex arena over two days and underwent bowl digging training where they learnt to retrieve a food reward buried under increasing amounts of sawdust as outlined in (Stuart et al., 2013) and described in further detail in **supplementary methods**. Following successful digging training rats underwent a single discrimination training session, where the rat was presented with two bowls filled with 2 cm of a different digging substrate e.g., sponge, perlite, cardboard squares etc. One of these substrates was consistently baited with one reward pellet (CS+) and the other was not rewarded (CS-). Both substrates were mixed with a crushed reward pellet to prevent odour discrimination. In the first trial, the rat explored both bowls until the pellet was found. In the second trial, as soon as the rat chose one bowl, the other was removed. If the rat chose the baited bowl, it was recorded as correct. This was repeated until the rat chose the baited bowl over 6 consecutive trials within a maximum of 20 trials. After this, the rat was considered fully trained and ready for testing.

### 2.10 Reward learning assay

The RLA is a 5 consecutive-day protocol consisting of 4 pairing sessions followed by a testing day as previously described by (Stuart et al., 2013). Each pairing session followed the same protocol as the discrimination session above (CS+ vs CS-). The rat learned two independent reward associations (CSA+ or CSB+), where one substrate was associated with one reward pellet and the other with two pellets. Over the course of the 4 pairing sessions, the rat is presented twice with substrate reward contingencies (CSA+ vs CS- or CSB++ vs CS-). On the testing day, the rat was presented with the two previously reinforced substrates (CSA vs CSB) (**fig 1**). The substrates were randomly reinforced with one reward pellet (probability 1 in 3 CSA or CSB) over the course of 30 trials. Each substrate was mixed with a crushed pellet to prevent odour discrimination. The position of the substrates, left or right, was pseudorandomised was . Substrate choice over the 30 trials and number of pellets consumed was recorded. % choice bias for the two-pellet paired substrate was the primary output measure and was calculated using (number of two-pellet paired choices/30*100)-50. A positive value indicates a reward -induced positive bias. Additional checks for substrate bias, directional bias and number of pellets consumed relative to chance (0 or 10) was performed.

**Fig.1.**
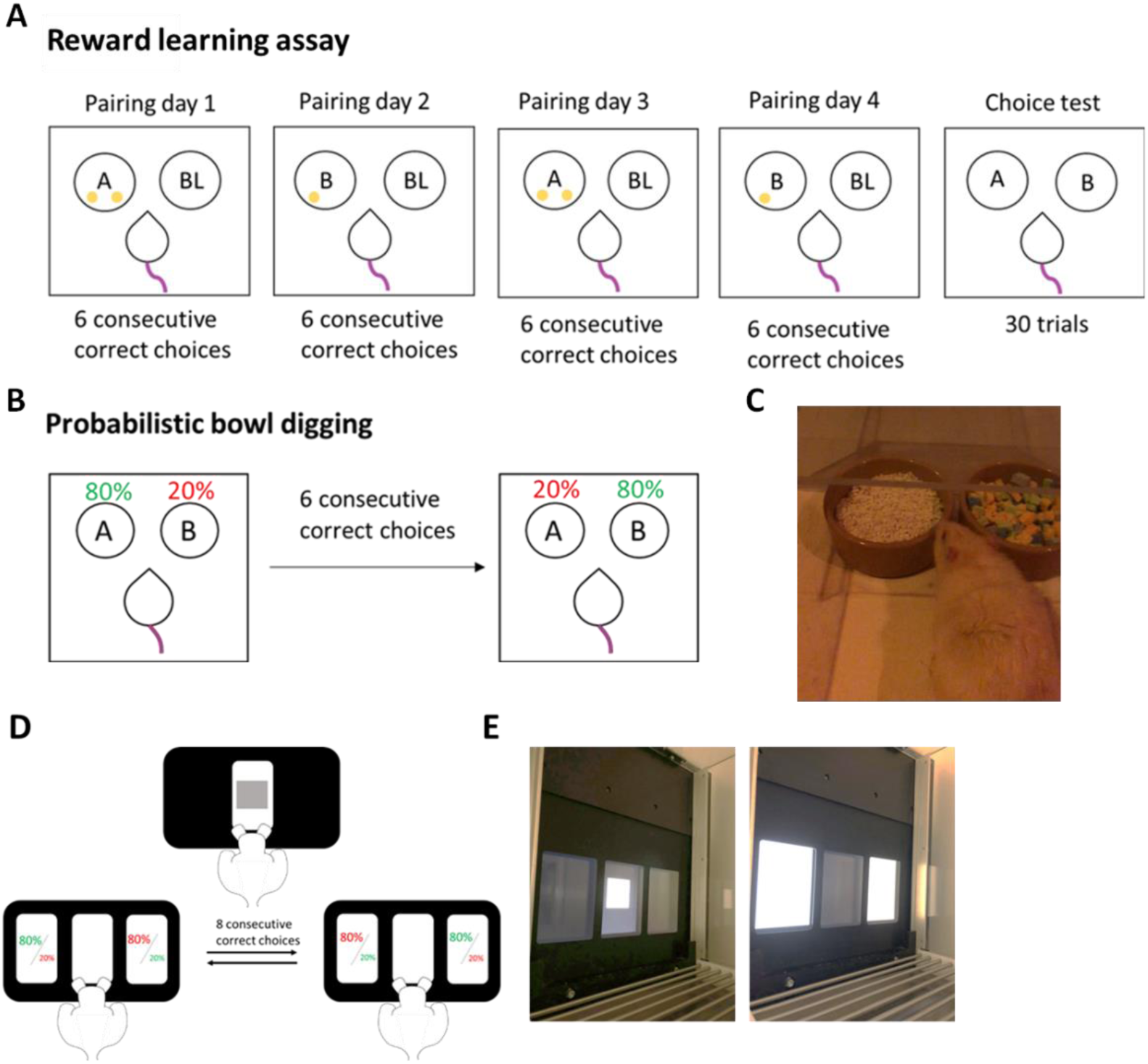
Reward learning and probabilistic reversal learning tasks. **A** The reward learning assay which consists of 4 pairing sessions followed by a choice test. **B** The probabilistic bowl digging task. **C** An aged rat making a digging decision. **D** The rat presses an initiation square and then must select either the left or right window, which are rewarded either 80% or 20 % of the time. **E** View from operant box. A = substrate A, B = substrate B, BL = blank substrate.

### 2.11 Probabilistic reversal learning bowl digging task

In a single session, the rat was presented with two bowls containing two different substrates. One substrate was baited with a single pellet 80 % of the time (‘rich’ substrate), and the other substrate baited 20 % of the time (‘lean’ substrate). The rat had to choose the correct (rich) bowl 6 times consecutively within a maximum of 30 trials to successfully ‘acquire’ the first rule. Once the rat had acquired this initial rule the contingencies were reversed such that the rich substrate became the lean substrate and *vice versa*. The rule was reversed every time the rat chose the rich bowl 6 times consecutively within 30 trials (**fig 1**). The position of the bowls was swapped pseudo randomly as described in the RLA. The session ended after a maximum of 1hr, or 4 consecutive omissions, or the rat failed to learn a rule within 30 trials. The primary output measures were number of reversals, win-stay probability and lose-stay probability, **summarised in table 2**.

**Table 2.** Key output measures for the probabilistic reversal learning task (bowl digging and touchscreen).

| Output measure | Description | Calculation |
| --- | --- | --- |
| **Reversals** | The number of times the rat learns the new rule following reversal of contingencies. |  |
| **Win-stay probability** | The probability that the rat will choose the bowl where it found a reward in the previous trial. | Win-stay probability = win-stay count / (win-stay count + win-shift count) |
| **Lose-shift probability** | The probability that the rat will shift its choice to the other bowl after not finding a reward in the previous trial. | Lose-shift probability = lose-shift count / (lose-shift count + lose-stay count) |

### 2.12 Probabilistic reversal learning operant task training

The operant box used for this task consisted of an infrared touchscreen panel with three windows where the animal could respond by touching them. A magazine was positioned centrally opposite the touchscreen panel where rats could access 45 mg reward pellets (TestDiet, Sandown Scientific, UK). The box also contained a house light and tone generator. The task was set up as previously described in (Wilkinson et al., 2020), which was based in the original design used in (Bari et al., 2010). Rats first underwent a 30 min session where a reward pellet was dispensed into the magazine automatically every 40 s. Rats then underwent continuous reinforcement training where touching the initiation square in the middle window resulted in the delivery of a reward pellet. This was accompanied by a tone and the magazine illuminating. The session finished after a maximum of 30 mins or the rat completed 120 trials. Once rats completed 120 trials over two consecutive days, they progressed to a second continuous reinforcement stage, where the rat was required to respond to the initiation square and then the illuminated window either to the left or right to receive a reward pellet. The session finished after a maximum of 40 mins or the rat completed 200 trials. If the rat touched the initiation square but took >10s to respond to the left or right panel, then a 10s timeout occurred where the house light came on and no reward was delivered. Once the rat had completed >120 trials over two consecutive sessions it was deemed trained and ready to undergo the probabilistic reversal learning task.

### 2.13 Probabilistic reversal learning task test sessions

One window was designated the ‘rich’ stimulus and was rewarded 80% of the time (delivery of reward pellet) and punished 20% of the time (10s timeout and house light on). The other window was designated the ‘lean’ stimulus and was punished 80% of the time and rewarded 20% of the time. The starting position of the rich stimulus was counterbalanced across cohort and remained consistent across sessions. When the rat made 8 consecutive rich choices, the contingencies swapped, such that the window that was previously designated rich became lean and vice versa (**fig.1**). The contingencies continued to change each time the new rich stimulus was selected 8 consecutive times up to a maximum of 40 mins or 200 trials. Successful learning of the first rule was termed ‘acquisition’ and successful learning of the new rule was termed a ‘reversal’.

### 2.14 Statistical analysis

In this study design, the single animal was considered the experimental unit. Data was tested for normality using the Shapiro-Wilk test. Where data was normally distributed, data were analysed with an independent t-test, or RM two-way ANOVA where performance across sessions or phases was additionally analysed. Where normality was violated, the Mann-Whitney U test was utilized. Significance was defined as p < 0.05. Where significant main effects or interactions were found these were reported in the results and followed up with appropriate post-hoc tests. Where a p value of <0.1 was found it was reported in the results as a trend level effect but not further analysed. *A priori* and consistent with other work (Marangoni et al., 2023), statistical outliers were defined as data points 2 standard deviations from the group mean. Where this occurred, these points were excluded, and replaced with the group mean to allow repeated measures analysis where required. If a rat had two outlying data points across sessions then the animal was excluded from the analysis for that measure. Instances where this occurred are reported in a table in **supplementary figures (S2)**. Statistical analysis and graphing were conducted using GraphPad Prism v 10.0 for windows.

## 3 Results

### 3. 1 Aged rats do not show impairment in physical effort-related choice behaviour

Rat performance in the progressive ratio task was analysed under food restricted and *ad libitum* feeding conditions. Under food restriction, not all rats reached a breakpoint and instead the last fixed ratio was used. There was no difference in last fixed ratio completed between age groups (p > 0.05) (**fig.2.A**). When only rats that reached a breakpoint were analysed, there was similarly no difference (p > 0.05) (**fig.2.B**). Under *ad libitum* feeding conditions, there was a trend towards aged rats having a lower breakpoint (t_(18)_ = 1.964, p = 0.0651) (**fig.2.C**).

**Figure 2.**
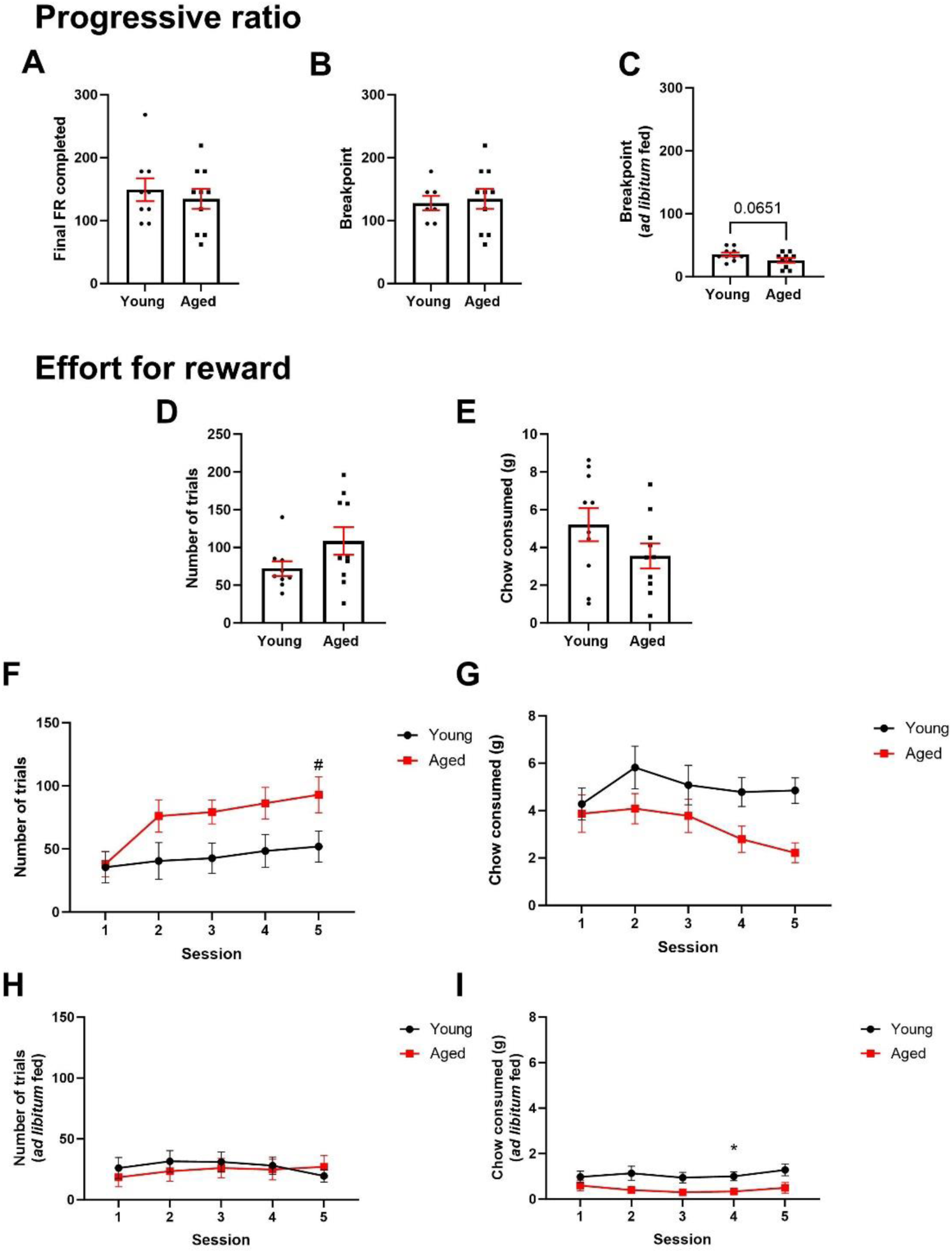
Aged rats do not show any motivational impairments in the progressive ratio or effort for reward tasks. **A** Under food restricted conditions, there was no difference in final ratio completed or **B** breakpoint in young versus aged rats (p > 0.05). **C** Under ad libitum feeding conditions there was a trend towards aged rats showing a reduced breakpoint. **D** In a single session of EfR directly after training, there was no difference in number of trials completed or **E** chow consumed (p > 0.05). **F** When repeated over 5 sessions, aged rats showed an increase in number of trials completed at day 5 vs day 1 (p < 0.05) **G** but no change in chow consumed (p > 0.05). Under ad libitum feeding conditions, there was no effect of age or session on number of trials (p > 0.05) **I** however aged rats consumed more chow on day 4 than younger rats (p < 0.05). *p< 0.05 (between-subject analysis) #p< 0.05 (within-subject analysis). Bars are mean ± SEM unless non-parametric where median ± interquartile range was used.

Rats were also tested in the effort for reward task, both directly after training in a single session, and over five consecutive sessions later in the behavioural battery. In the single session, there was no difference in number of trials completed or chow consumed (p > 0.05) (**fig.2.D&E**). When repeated over five sessions, there was a session*age interaction (F_(4,64)_ = 4.070, p = 0053). There was also a main effect of session (F_(2.126, 34.01)_ = 12.01, p < 0.0001) and a trend towards a main effect of age (F_(1,16)_ = 3.643, p = 0.0744). Post-hoc analysis revealed that number of trials completed by aged rats increased across sessions compared to session 1 (p = 0.0099 - 0.0162). Post hoc analysis of age comparison across sessions did not reach significance (p > 0.05) (**fig.2.F**). Analysis of chow consumed across sessions revealed a main effect of session (F_(2.587, 46.57)_ = 3.0, p = 0.0469. However, post hoc analysis did not reach significance. There was also a trend towards a main effect of age (F_(1,18)_ = 4.375, p = 0.0509) (**fig.2.G**). Under *ad libitum* feeding conditions, there was no effect of session or age on number of trials (P > 0.05) (**fig.2.H**). However, there was a main effect of age on chow consumed (F_(1,18)_ = 7.178, p = 0.0153). Post-hoc analysis revealed that younger rats ate more chow than aged rats on session 4 (**fig.2.I**).

### 3.2 Aged rats do not show behaviours or physiological changes consistent with emotional blunting

Young and aged rats underwent a range of tests relating to emotional reactivity. In the novelty suppressed feeding test, there was no difference in latency to approach the bowl of chow (p > 0.05), but aged rats took longer to eat from the bowl than younger rats (t_(18)_ = 2.664, p = 0.0158) (**fig.3.A&B**). In a consumption test, aged rats consumed more lab chow within 10 minutes than younger rats (t_(18)_ = 2.416, p = 0.0266) (**fig.3.C**). Analysis of blood plasma corticosterone levels before and after restraint stress revealed a main effect of stress (F_(1,17)_ = 60.74, p < 0.0001) but no effect of age or stress*age interaction. Post-hoc analysis showed that plasma corticosterone was higher following stress in both young and aged rats (p = 0.0001 and p < 0.0001 respectively) (**fig.3.D**).

**Figure 3.**
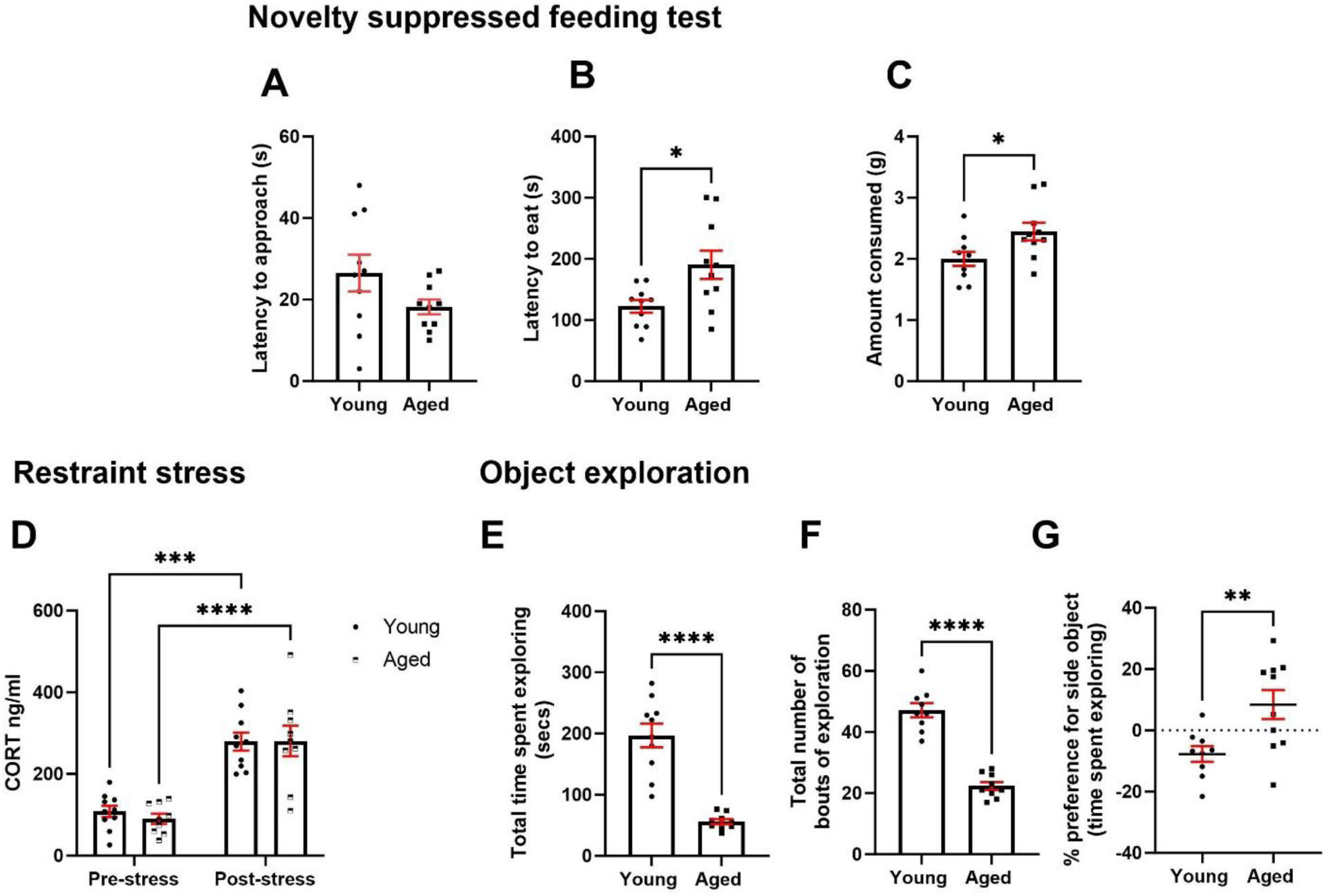
Aged rats do not show behaviours or physiological changes consistent with emotional blunting. **A** There was no age difference in latency to approach the centrally placed bowl in the NSFT p > 0.05. **B** Aged rats took longer to eat from the bowl (p < 0.05). **C** Aged rats consumed more chow in a 10 min consumption test than young rats (p < 0.05). **D** Both young and aged rats showed an increase in blood plasma CORT levels following restraint stress. **E** Aged rats spent less time and **F** bouts of exploration of novel objects than young rats. **G** Aged rats showed a greater preference for the side placed object compared to younger rats (p < 0.05). *p< 0.05, **p<0.01, ****p<0.0001. Bars are mean ± SEM unless non-parametric where median ± interquartile range was used.

In a position-based exploration test, aged rats spent less time and had less bouts of exploring objects (t_(17)_ = 6.789, p < 0.0001 and t_(16)_ = 9.357, p < 0.0001) (**fig.3.E&F**). In addition, aged rats showed a greater preference for exploring an object placed at the side versus in the middle of an open field arena (t_(17)_

= 2.192, p = 0.0097) (**fig.3.G**).

### 3.3 Aged rats show intact reward-related cognition and reward sensitivity but impaired cognitive flexibility in a bowl-digging version of the probabilistic reversal learning task

In the sucrose preference test, while both young and aged rats showed a preference for 2 % sucrose solution over standard tap water (p < 0.001), aged rats showed a greater % sucrose preference than younger rats (t_(15)_ = 2.191, p = 0.0447) (**fig.4.A**). In the reward learning assay, both young and aged rats showed a preference for the two-pellet paired substrate (t_(8)_ = 10.09, p < 0.0001 and t_(7.222)_ = 7.0, p = 0.0002 respectively) but there was no difference between groups (p > 0.05) (**fig.4.B**). In the probabilistic reversal learning bowl digging assay, there was a trend towards aged rats taking more trials to learn the first rule (t_(14)_ = 1.876, p = 0.0817) (**fig.4.C**). Aged rats completed less reversals than young rats (t_(14)_ = 5.001, p = 0.0002) (**fig.4.D**). Analysis of win-stay probability showed a main effect of age (F_(1,17)_ = 12.21, p = 0.0028), a main effect of phase (F_(1,14)_ = 24.42, p = 0.0002) and an age*phase interaction (F_(1,14)_ = 10.56, p = 0.0058). Post-hoc analysis revealed that young rats showed a higher win-stay probability in the reversal phase compared to aged rats (p = 0.0002). In addition, aged rats showed a reduction in win-stay probability in the reversal phase versus the acquisition phase (p = 0.0002) (**fig.4.E**). There was no effect of age or phase on lose-shift behaviour (p > 0.05) (**fig.4.F**).

**Figure 4.**
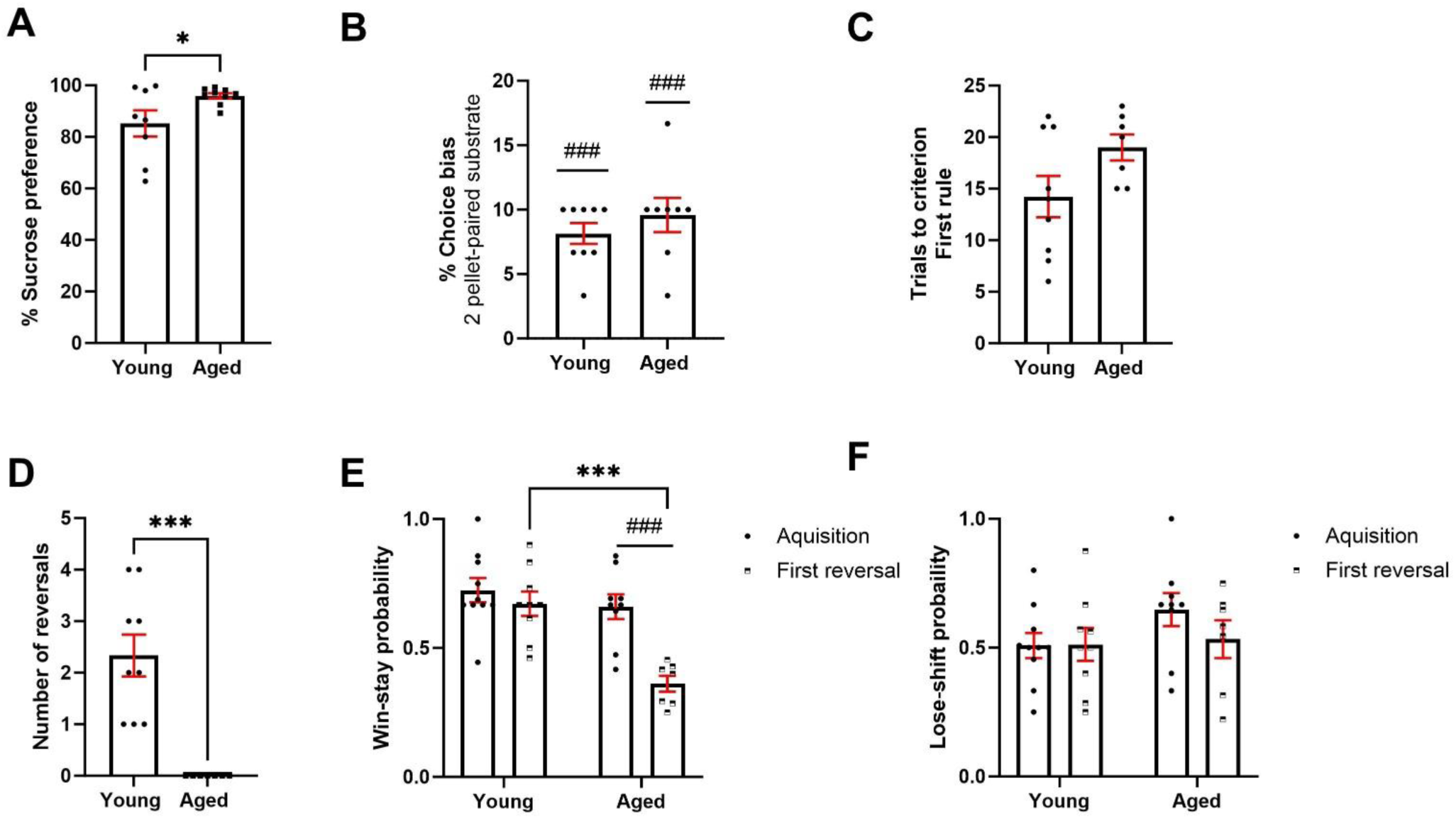
Aged rats show intact reward-related cognition and reward sensitivity but impaired cognitive flexibility in a bowl-digging version of the PRLT. **A** Both groups show a preference for sucrose, but aged rats show a greater preference than younger rats (p < 0.05). **B** Both groups showed a preference for the 2 pellet paired substrate in the RLA (p < 0.001) but no difference between age groups. **C** There was no difference in number of trials taken to learn first rule in the PRLT (p > 0.05). **D** Aged rats reversed less than younger rats in the PRLT (p < 0.001). **E** In the acquisition phase, aged rats had a lower win-stay probability than young rats (p < 0.001) and also had a lower win-stay probability in the reversal versus acquisition phase (p < 0.001). **F** There was no effect of age or phase on lose-shift behaviour (p > 0.05). *p< 0.05 (between-subject analysis) #p< 0.05 (within-subject analysis). Bars are mean ± SEM unless non-parametric where median ± interquartile range was used.

### 3.4 Aged rats show no impairment in an operant touchscreen version of the probabilistic reversal learning task

There was no effect of age or session on the trial at which the first trial was learned (p > 0.05) (**fig.5.A**). However, there was a main effect of session (F_(3.251, 55.81)_ = 5.719, p = 0.0013) and a session*age interaction (F_(6, 103)_ = 2.803, p = 0.0137) on number of reversals. However post-hoc analysis revealed no differences across session or age (p > 0.05) (**fig.5.B**). There was also a main effect of session on win-stay probability (F_(3.454, 62.17)_ = 4.219, p = 0.0064) but no effect of age (p > 0.05). Post-hoc analysis revealed that win-stay probability was increased at session 6 and 7 compared to session 1 in the young group (p = 0.013 and p = 0.04 respectively) (**fig.5.C**). There was a main effect of session on lose-shift probability (F_(3.07, 52.19)_ = 3.152, p = 0.0315) but no effect of age. However, post-hoc analysis revealed no difference in session performance compared to session 1 (**fig.5.D**). Finally, there was a main effect of session on initiation time (F_(2.853, 45.65)_ = 5.448, p = 0.0032). Post-hoc analysis revealed young rats showed a decrease in initiation time at day 6 and day 7 compared to day 1 (p = 0.00253 and p = 0.0167 respectively). There was no effect of age or age*session interaction (p > 0.05) (**fig.5.E**).

**Figure 5.**
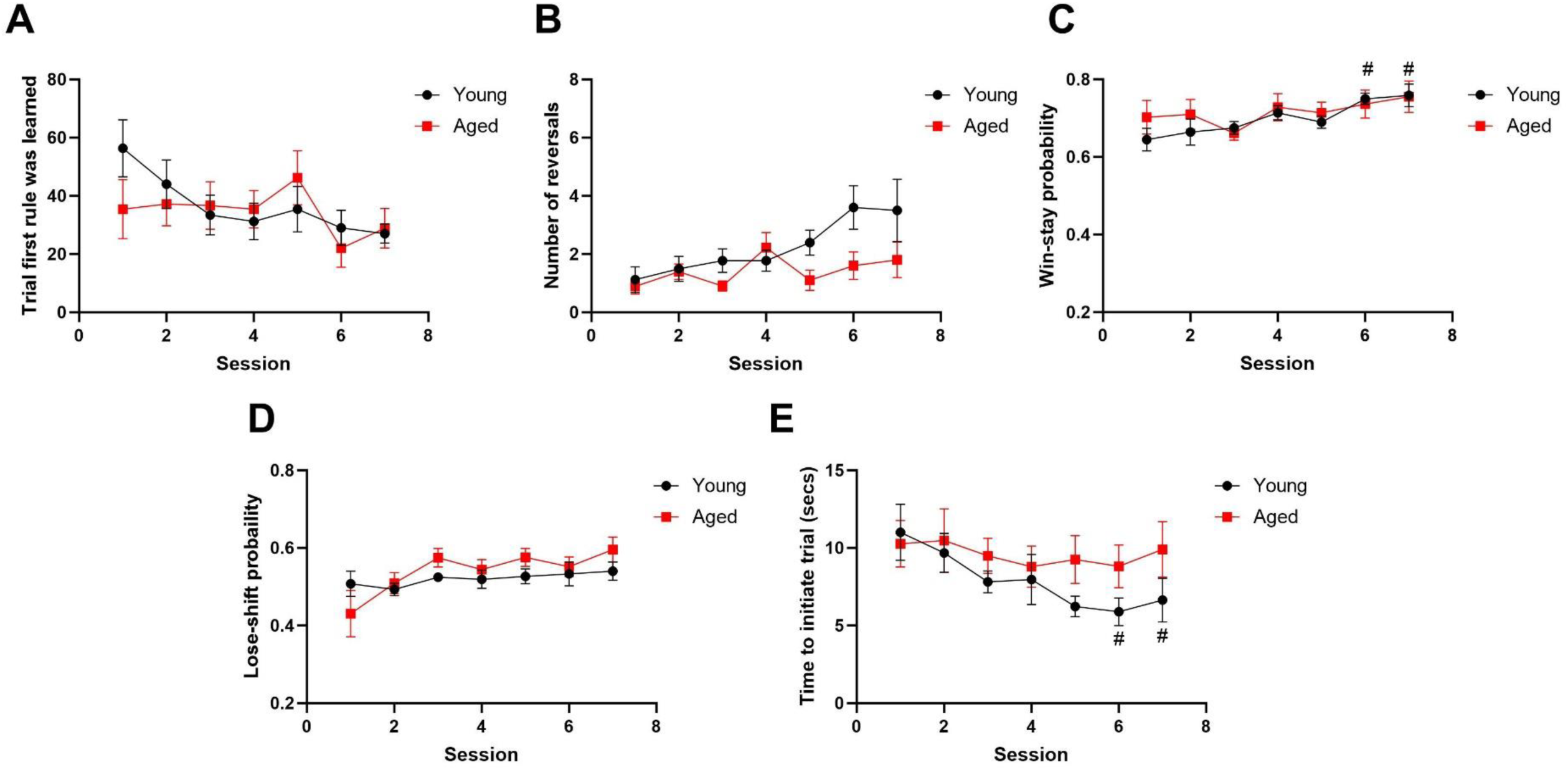
There was no effect of age on cognitive flexibility in the operant version of the PRLT. **A** There was no post-hoc effect of age on trial first rule was learned or **B** number of reversals (p > 0.05). **C** There was no effect of age on win-stay probability however win-stay probability was increased in the young group at session 6 and 7 versus 1 (p < 0.05). **D** There was no post-hoc level effect of age or session on lose-shift probability. **E** There was no effect of age on trial initiation time, however young rats showed a lower initiation time at session 6 and 7 versus 1. #p<0.05 (within-subject comparison). Points are mean ± SEM.

A summary of behavioural results and their comparison with previously published behavioural data from aged mice is provided in **table 3**.

**Table 3.** Summary of behavioural changes in the aged animal versus the young animal. NC = no change. NA = not determined, test was not conducted in that species. Blue-behaviours relating to behaviour-cognition domain (motivation), peach-behaviours relating to emotional domain, yellow-behaviours relating to behaviour-cognition domain (cognition).

| Behavioural task/physiological measure | Rat | Mouse (Jackson et al., 2021) |
| --- | --- | --- |
| Position-based object exploration: total exploration | Reduced | ND |
| Position-based object exploration: position bias | Increased | ND |
| Novelty suppressed feeding test | Increased latency to feed | Decreased latency to feed |
| Effort for reward | NC | Decreased trials and chow consumed |
| Progressive ratio | NC | Decreased final ratio achieved |
| Hormonal response to restraint stress | NC | Decreased levels |
| Reward learning assay | NC | ND |
| Sucrose preference test | Increased preference | Decreased preference |
| Probabilistic reversal learning (bowl digging) | Decreased reversals | ND |
| Probabilistic reversal learning (touchscreen) | NC | ND |

## 4 Discussion

In this study we used a battery of behavioural tests hypothesised to map onto different behavioural dimensions of apathy syndrome, including motivational, emotional and cognitive components. Use of two operant effort-based decision making tasks found a trend towards an increase in physical effort-based responding in aged rats but no overall difference. However aged rats did show a reduction in spontaneous object exploration. Aged rats showed a greater latency to consume food in the novelty suppressed feeding task (NSFT) and preference for exploration of the side versus middle object in a position-based exploration test consistent with a higher level of anxiety-like behaviour rather than emotional blunting. Aged rats showed a corticosterone response to a restraint stressor equivalent to younger rats suggesting increased emotional reactivity is restricted to the behavioural level. Aged rats showed a profound impairment in reversal behaviour in a bowl digging version of the probabilistic reversal learning test. The impairment appeared specific to a loss in win-stay but not lose-stay behaviour and suggests perseverative behaviour. This impairment was not driven by changes to core affective state or reward-related cognition (reward learning assay (RLA)) or changes in reward sensitivity (sucrose preference test (SPT)). Of note, the operant version of this task did not pick up these same differences. Here, we consider how these behavioural findings relate to the psychiatric syndrome apathy and findings in aged mice (Jackson et al., 2021) and consider the overall suitability of the use of aged rats as a model of apathy.

### 4.1 Aged rats show intact physical effort-based decision making

Patients with apathy exhibit a deficit in motivation, including some disruption to effort-based decision making (Le Heron et al., 2018). The effort for reward (EfR) task and progressive ratio (PR) task are considered translational measures of effort based decision making (Salamone et al., 1991, Treadway et al., 2009, Stoops, 2008, Richardson and Roberts, 1996). These tasks have been shown to be sensitive to a range of manipulations that modulate motivational state in animals, including ageing in mice where a reduction in high effort, high value reward responding and an increase in low effort, low value reward engagement was found (Jackson et al., 2021) . Deficits have also been found in patients with apathy associated with Huntington’s Disease (Heath et al., 2019) and neurocognitive disorders (Zeghari et al., 2021). We found no evidence of impairment in aged rats, and in fact, there was a trend towards an increase in high effort choices in the aged rats, suggesting effort-based decision making is not affected by ageing in this species. This may in part be driven by behavioural context. It has previously been shown that, in contrast to other effort-based behaviours, instrumental lever press responding is conserved in aged rats (Samson et al., 2014). Therefore, in addition to assessing motivated behaviour in a conditioned responding context, we assessed motivation to explore novel objects. Previous work has shown that exploration of novelty is a motivated behaviour based on an intrinsic drive to forage and seek reward. Novelty-based motivation has been found to be sensitive to ageing at in humans and rodents (Düzel et al., 2010) and has also been shown to be reduced in patients with apathy (Yamagata et al., 2004, Kaufman et al., 2016). We found that aged rats exhibit a robust reduction in exploration of novel objects, suggestive of a motivational impairment in this context. Thus, operant-based conditioning methods may be less sensitive to motivational deficits in this species and more ethologically relevant, forage-based methods may provide a more sensitive readout of motivational state in rats.

### 4.2 Aged rats show heightened affective reactivity inconsistent with emotional blunting

Patients with apathy often exhibit emotional blunting symptoms, where they show limited reactivity to environmental stimuli (Levy and Dubois, 2006). In animal models, this can be quantified by assessment of behavioural and physiological response to behaviours relating to the negative valence domain (Jackson and Robinson, 2022, Jackson et al., 2021), where a reduction in appropriate response is indicative of affective blunting. The NSFT is a commonly-used measure of behaviours relating to anxiety, where the rodent must overcome its aversion of a novel, open space to eat from a bowl placed in the center (Bodnoff et al., 1988). We found that aged rats took longer to eat from a centrally placed bowl, which is indicative of heightened anxiety-like behaviour (Bodnoff et al., 1988). Age-related metabolic differences have the potential to confound appetitive tasks. We found that aged rats consumed more lab chow in a consumption test and therefore loss of appetite does not explain increased latency to consume food. However we cannot fully rule out age-related metabolic differences driving at least part of this behaviour. Therefore, to assess anxiety-like behaviour in a non-appetitive context, we developed a position-based exploration test based on the well-characterized principle that a rodent will exhibit thigmotaxis when placed in an open arena (Hall and Ballachey, 1932). As such, the rodent will bias its exploration of an object positioned to the side of the arena versus the object placed in the center. A stronger exploration bias may indicate a greater level of anxiety-related behaviour. Aged rats showed a stronger bias for exploration of the object placed at the side rather than the center of the arena. Thus, consistent with the results from the NSFT, aged rats show increased anxiety-like behaviour. Aged rats exhibited a similar increased corticosterone response to restraint stress to that of younger rats, suggesting this increase in reactivity is conserved to the behavioural level. These findings contrast with our prediction of a blunted response and our findings in aged mice, where a blunted restraint-induced corticosterone response was observed and reduced anxiety-related behaviour in the NSFT, indicative of emotional blunting (Oh et al., 2018, Jackson et al., 2021). These findings further highlight key species differences in behaviours relevant to apathy with no evidence of emotional blunting observed in aged rats but an increase in anxiety-related behaviours.

### 4.3 Aged rats show a reduction in cognitive flexibility but only in a bowl-digging-based task

Impairments in cognitive flexibility have been found across conditions where apathy commonly occurs (Nilsson et al., 2015). The rodent PRLT is usually conducted as a touchscreen assay (Der-Avakian et al., 2013, Wilkinson et al., 2020, Bryce and Howland, 2015), which is characterised by extended training periods, large numbers of trials and repeated sessions which may result in performance becoming more dependent on procedural learning strategies. Bowl digging assays may provide more ecologically relevant learning and have been used previously to assess affective state, attention and cognitive flexibility (Stuart et al., 2013, Birrell and Brown, 2000, McAlonan and Brown, 2003). Here, we found most rats successfully learned to select the rewarded substrate and while there was no group difference, there was a trend towards aged rats taking more trials to learn in the acquisition phase. Strikingly, when the rule was reversed, all aged rats were unable to reverse, consistent with an impairment in cognitive flexibility. Assessment of likelihood that the rat would choose the previously rewarded bowl (win-stay probability) showed that while win-stay behaviour was equivalent to younger rats in the acquisition phase, it was markedly reduced in the reversal phase. This deficit was not seen in lose-shift behaviour, the likelihood the rat will choose the other substrate on the next trial after it was unrewarded on the previous trial. Therefore, this reversal learning impairment is likely due to a failure to remove prior associations with a positive cue, known as ‘perseverative’ behaviour (Nilsson et al., 2015). Perseverative behaviour in old age has been widely reported in other rodent studies. Aged rats could learn normally but showed impaired reversal in a spatial T maze discrimination task, odour discrimination task and visual discrimination task, showing this behaviour is consistent over multiple sensory domains (Stephens et al., 1985, Brushfield et al., 2008). Elderly humans also show impairments in reversal learning and a mild impairment in acquisition learning compared to younger adults (Weiler et al., 2008).

These changes were not driven by differences in core affective state or reward-related cognition, as revealed by the RLA, where both young and aged rats showed a robust and equivalent reward-induced positive bias. This task has been shown to be a sensitive measure of reward-related cognition and core affective state changes, where models such as early life adversity, chronic pain and chronic administration of pro-depressant drugs impairs reward learning (Stuart et al., 2019, Stuart et al., 2017)(Phelps et al., 2021). These changes were also not driven by differences in reward sensitivity, as aged rats showed an intact and even enhanced preference for 2% sucrose solution over standard tap water in the SPT. Thus, this appears to be an impairment specific to cognitive flexibility.

Of note, we do not observe this same deficit either in terms of absolute reversals or win-stay probability in the touchscreen version of the PRLT. Here, we observed similar levels of performance between groups across sessions. Therefore, the operant version of the PRLT may be less sensitive to age-related differences in cognitive flexibility.

It is important to note that the use of aged females would improve overall translatability of findings, though they were not made available to us due to increased risk of age-related impairments. Due to limited data availability, it is unclear whether aged females would show a similar pattern of deficits in behaviours relating to apathy however there is some limited evidence for differences in motivated behaviour (van Hest et al., 1988). Differences in lifespan between the rat and mouse make it difficult to make direct comparisons between species (Sengupta, 2013, The Jackson Laboratory, 2021). While both species were tested in a window considered to be ‘aged’, the length of the aged phase may differ between species. This may induce variation in when age-related impairments develop. It is also not known the extent to which the laboratory environment and associated breeding has impacted on ageing in these species.

## 5 Conclusion

We found limited evidence for effort-based motivational deficit or emotional blunting in the rat but observed an increase in anxiety-related behaviours and impairments in cognitive flexibility in a bowl-digging reversal learning task. The findings in aged rats were very different from those observed in aged mice across these different behavioural domains suggesting there are key species difference in apathy-relevant behavioural changes associated with ageing. The evidence for a reduction in cognitive flexibility and perseverative behaviour in aged rats may be useful in relation to understanding more about age-related cognitive flexibility and the use of a more ethologically relevant task seems to increase sensitivity to these impairments when compared to a touchscreen task with prolonged training periods and repeated exposure to both acquisition and reversal learning phases. However, this does not address the challenge of studying apathy based on impairments across multiple domains. The behavioural impairments observed in mice suggested both motivational impairments and emotional blunting develop in ageing, but evidence of cognitive impairments in a novel object task was not observed. It should be noted that this task measures recognition memory and is not sensitive to changes in cognitive flexibility. In contrast, aged rats showed either no change or an increase in motivation in effort-related decision-making tasks which were not related to appetite. They also exhibited an increase in anxiety-related behaviours. Overall, these studies do not support the use of an aged rat model (21-27 months) to study apathy as a syndrome however, aged mice may represent a more translationally relevant model and further evaluation of their cognitive flexibility would be useful. It might also be interesting to investigate further the species differences seen between rats and mice and expand the behavioural characterization of ageing in these two important species.

## Acknowledgements

The authors gratefully acknowledge the technical support of Mrs Julia Bartlett and the provision of analysis code for the probabilistic reversal learning task from Dr Matthew Wilkinson. We also acknowledge the support of Dr Hugh Marston and the provision of aged rats from Eli Lilly & co.

## Data availability

All data used in this article are available and can be accessed via the Open Science Framework upon publication.

## Abbreviations

CORT: corticosterone
CS: conditioned stimulus
EfR: effort for reward
FR: fixed ratio
NSFT: novelty supressed feeding test
PR: progressive ratio
PRLT: probabilistic reversal learning task
RIA: radioimmunoassay
RLA: reward learning assay
RM: repeated measures
SPT: sucrose preference test

## References

Ang, Y.-S., Lockwood, P., Apps, M. A. J., Muhammed, K. & Husain, M. 2017. Distinct Subtypes of Apathy Revealed by the Apathy Motivation Index. PLoS ONE, 12, e0169938.10.1371/journal.pone.0169938

Bari, A., Theobald, D. E., Caprioli, D., Mar, A. C., Aidoo-Micah, A., Dalley, J. W. & Robbins, T. W. 2010. Serotonin Modulates Sensitivity to Reward and Negative Feedback in a Probabilistic Reversal Learning Task in Rats. Neuropsychopharmacology, 35, 1290–1301.10.1038/npp.2009.233

Beck, A. 1967. Depression: Clinical, experimental and theoretical aspects London. Staples Press.

Bodnoff, S. R., Suranyi-Cadotte, B., Aitken, D. H., Quirion, R. & Meaney, M. J. 1988. The effects of chronic antidepressant treatment in an animal model of anxiety. Psychopharmacology, 95, 298–302.10.1007/BF00181937

Brodaty, H., Altendorf, A., Withall, A. & Sachdev, P. 2009. Do people become more apathetic as they grow older? A longitudinal study in healthy individuals. International Psychogeriatrics, 22, 426–436.10.1017/S1041610209991335

Bryce, C. A. & Howland, J. G. 2015. Stress facilitates late reversal learning using a touchscreen-based visual discrimination procedure in male Long Evans rats. Behavioural Brain Research, 278, 21–28.10.1016/j.bbr.2014.09.027

Cuthbert, B. N. 2015. Research Domain Criteria: toward future psychiatric nosologies. Dialogues in clinical neuroscience, 17, 89–97.10.31887/DCNS.2015.17.1/bcuthbert

Der-Avakian, A., D’souza, M. S., Pizzagalli, D. A. & Markou, A. 2013. Assessment of reward responsiveness in the response bias probabilistic reward task in rats: implications for cross-species translational research. Transl Psychiatry, 3, e297.10.1038/tp.2013.74

Düzel, E., Bunzeck, N., Guitart-Masip, M. & DÜzel, S. 2010. NOvelty-related Motivation of Anticipation and exploration by Dopamine (NOMAD): Implications for healthy aging. Neuroscience & Biobehavioral Reviews, 34, 660–669.10.1016/j.neubiorev.2009.08.006

Gerritsen, D. L., Jongenelis, K., Steverink, N., Ooms, M. E. & Ribbe, M. W. 2005. Down and drowsy? Do apathetic nursing home residents experience low quality of life? Aging & Mental Health, 9, 135–141.10.1080/13607860412331336797

Griesius, S., Mellor, J. R. & Robinson, E. S. J. 2020. Comparison of acute treatment with delayed-onset versus rapid-acting antidepressants on effort-related choice behaviour. Psychopharmacology, 237, 2381–2394.10.1007/s00213-020-05541-9

Hall, C. & Ballachey, E. L. 1932. A study of the rat’s behavior in a field. A contribution to method in comparative psychology. University of California Publications in Psychology, 6, 1–12

Heath, C. J., O’callaghan, C., Mason, S. L., Phillips, B. U., Saksida, L. M., Robbins, T. W., Barker, R. A., Bussey, T. J. & Sahakian, B. J. 2019. A Touchscreen Motivation Assessment Evaluated in Huntington’s Disease Patients and R6/1 Model Mice. Frontiers in Neurology, 10.10.3389/fneur.2019.00858

Ishii, S., Weintraub, N. & Mervis, J. R. 2009. Apathy: A Common Psychiatric Syndrome in the Elderly. Journal of the American Medical Directors Association, 10, 381–393.10.1016/j.jamda.2009.03.007

Jackson, M. G., Lightman, S. L., Gilmour, G., Marston, H. & Robinson, E. S. J. 2021. Evidence for deficits in behavioural and physiological responses in aged mice relevant to the psychiatric symptom of apathy. Brain Neurosci Adv, 5, 23982128211015110.10.1177/23982128211015110

Jackson, M. G. & Robinson, E. S. J. 2022. The importance of a multidimensional approach to the preclinical study of major depressive disorder and apathy. Emerg Top Life Sci, 6, 479–489.10.1042/etls20220004

Le Heron, C., Plant, O., Manohar, S., Ang, Y.-S., Jackson, M., Lennox, G., Hu, M. T. & Husain, M. 2018. Distinct effects of apathy and dopamine on effort-based decision-making in Parkinson’s disease. Brain 141, 1455–1469.10.1093/brain/awy110

Levy, R. & Dubois, B. 2006. Apathy and the Functional Anatomy of the Prefrontal Cortex–Basal Ganglia Circuits. Cerebral Cortex, 16, 916–928.10.1093/cercor/bhj043

Malpetti, M., Jones, P. S., Tsvetanov, K. A., Rittman, T., Van Swieten, J. C., Borroni, B., Sanchez-Valle, R., Moreno, F., Laforce, R., Graff, C., Synofzik, M., Galimberti, D., Masellis, M., Tartaglia, M. C., Finger, E., Vandenberghe, R., De Mendonça, A., Tagliavini, F., Santana, I., Ducharme, S., Butler, C. R., Gerhard, A., Levin, J., Danek, A., Otto, M., Frisoni, G. B., Ghidoni, R., Sorbi, S., Heller, C., Todd, E. G., Bocchetta, M., Cash, D. M., Convery, R. S., Peakman, G., Moore, K. M., Rohrer, J. D., Kievit, R. A., Rowe, J. B. & Initiative, G. F. T. D. 2021. Apathy in presymptomatic genetic frontotemporal dementia predicts cognitive decline and is driven by structural brain changes. Alzheimer’s & Dementia, 17, 969–983.10.1002/alz.12252

Marangoni, C., Tam, M., Robinson, E. S. J. & Jackson, M. G. 2023. Pharmacological characterisation of the effort for reward task as a measure of motivation for reward in male mice. Psychopharmacology.10.1007/s00213-023-06420-9

Metha, J. A., Brian, M. L., Oberrauch, S., Barnes, S. A., Featherby, T. J., Bossaerts, P., Murawski, C., Hoyer, D. & Jacobson, L. H. 2020. Separating Probability and Reversal Learning in a Novel Probabilistic Reversal Learning Task for Mice. Frontiers in Behavioral Neuroscience, 13.10.3389/fnbeh.2019.00270

Nilsson, S. R. O., Alsiö, J., Somerville, E. M. & Clifton, P. G. 2015. The rat’s not for turning: Dissociating the psychological components of cognitive inflexibility. Neuroscience and biobehavioral reviews, 56, 1–14.10.1016/j.neubiorev.2015.06.015

Oh, H.-J., Song, M., Kim, Y. K., Bae, J. R., Cha, S.-Y., Bae, J. Y., Kim, Y., You, M., Lee, Y., Shim, J. & Maeng, S. 2018. Age-Related Decrease in Stress Responsiveness and Proactive Coping in Male Mice. Frontiers in aging neuroscience, 10, 128–128.10.3389/fnagi.2018.00128

Padala, P. R., Desouza, C. V., Almeida, S., Shivaswamy, V., Ariyarathna, K., Rouse, L., Burke, W. J. & Petty, F. 2008. The impact of apathy on glycemic control in diabetes: a cross-sectional study. Diabetes Res Clin Pract, 79, 37–41.10.1016/j.diabres.2007.06.012

Peterson, D. A., Elliott, C., Song, D. D., Makeig, S., Sejnowski, T. J. & Poizner, H. 2009. Probabilistic reversal learning is impaired in Parkinson’s disease. Neuroscience, 163, 1092–1101.10.1016/j.neuroscience.2009.07.033

Phelps, C. E., Lumb, B. M., Donaldson, L. F. & Robinson, E. S. 2021. The partial saphenous nerve injury model of pain impairs reward-related learning but not reward sensitivity or motivation. Pain, 162, 956–966.10.1097/j.pain.0000000000002177

Richardson, N. R. & Roberts, D. C. S. 1996. Progressive ratio schedules in drug self-administration studies in rats: a method to evaluate reinforcing efficacy. Journal of Neuroscience Methods, 66, 1–11.10.1016/0165-0270(95)00153-0

Robert, P., Lanctôt, K. L., Agüera-Ortiz, L., Aalten, P., Bremond, F., Defrancesco, M., Hanon, C., David, R., Dubois, B., Dujardin, K., Husain, M., König, A., Levy, R., Mantua, V., Meulien, D., Miller, D., Moebius, H. J., Rasmussen, J., Robert, G., Ruthirakuhan, M., Stella, F., Yesavage, J., Zeghari, R. & Manera, V. 2018. Is it time to revise the diagnostic criteria for apathy in brain disorders? The 2018 international consensus group. European Psychiatry, 54, 71–76.10.1016/j.eurpsy.2018.07.008

Roberts, D. C. S. & Richardson, N. R. 1992. Self-Administration of Psychomotor Stimulants Using Progressive Ratio Schedules of Reinforcement. In: Boulton, A. A., Baker, G. B. & Wu, P. H. (eds.) Animal Models of Drug Addiction. Totowa, NJ: Humana Press.10.1385/0-89603-217-5:233.

Salamone, J. D., Steinpreis, R. E., Mccullough, L. D., Smith, P., Grebel, D. & Mahan, K. 1991. Haloperidol and nucleus accumbens dopamine depletion suppress lever pressing for food but increase free food consumption in a novel food choice procedure. Psychopharmacology, 104, 515–521.10.1007/BF02245659

Samson, R. D., Venkatesh, A., Patel, D. H., Lipa, P. & Barnes, C. A. 2014. Enhanced performance of aged rats in contingency degradation and instrumental extinction tasks. Behavioral neuroscience, 128, 122–133.10.1037/a0035986

Sengupta, P. 2013. The Laboratory Rat: Relating Its Age With Human’s. International journal of preventive medicine, 4, 624–630

Stoops, W. W. 2008. Reinforcing effects of stimulants in humans: sensitivity of progressive-ratio schedules. Experimental and clinical psychopharmacology, 16, 503–512.10.1037/a0013657

Stuart, S. A., Butler, P., Munafò, M. R., Nutt, D. J. & Robinson, E. S. J. 2013. A Translational Rodent Assay of Affective Biases in Depression and Antidepressant Therapy. Neuropsychopharmacology, 38, 1625–1635.10.1038/npp.2013.69

THE JACKSON LABORATORY. 2021. WHEN ARE MICE CONSIDERED OLD? [Online]. Available: https://www.jax.org/news-and-insights/jax-blog/2017/november/when-are-mice-considered-old# [Accessed 19/05/2021 2021].

Treadway, M. T., Buckholtz, J. W., Schwartzman, A. N., Lambert, W. E. & Zald, D. H. 2009. Worth the ‘EEfRT’? The Effort Expenditure for Rewards Task as an Objective Measure of Motivation and Anhedonia. PLOS ONE, 4, e6598.10.1371/journal.pone.0006598

Van Hest, A., Van Haaren, F. & Van De Poll, N. E. 1988. The behavior of male and female Wistar rats pressing a lever for food is not affected by sex differences in food motivation. Behavioural Brain Research, 27, 215–221.10.1016/0166-4328(88)90118-0

Wilkinson, M. P., Grogan, J. P., Mellor, J. R. & Robinson, E. S. J. 2020. Comparison of conventional and rapid-acting antidepressants in a rodent probabilistic reversal learning task. Brain and Neuroscience Advances, 4.10.1177/2398212820907177

Windle, R. J., Wood, S. A., Shanks, N., Lightman, S. L. & Ingram, C. D. 1998. Ultradian Rhythm of Basal Corticosterone Release in the Female Rat: Dynamic Interaction with the Response to Acute Stress*. Endocrinology, 139, 443–450.10.1210/endo.139.2.5721

Zeghari, R., Lockwood, P. L., L’yvonnet, T., Jo, C., Robert, P. & Manera, V. 2021. TaPiscine: An effort-based decision-making task for apathy assessment in people with neurocognitive disorders. Alzheimer’s & Dementia, 17, e057746.10.1002/alz.057746

