## Supplementary figures for "Characterisation of behaviours relevant to apathy syndrome in the aged male rat"

Supplementary figure

**S1**
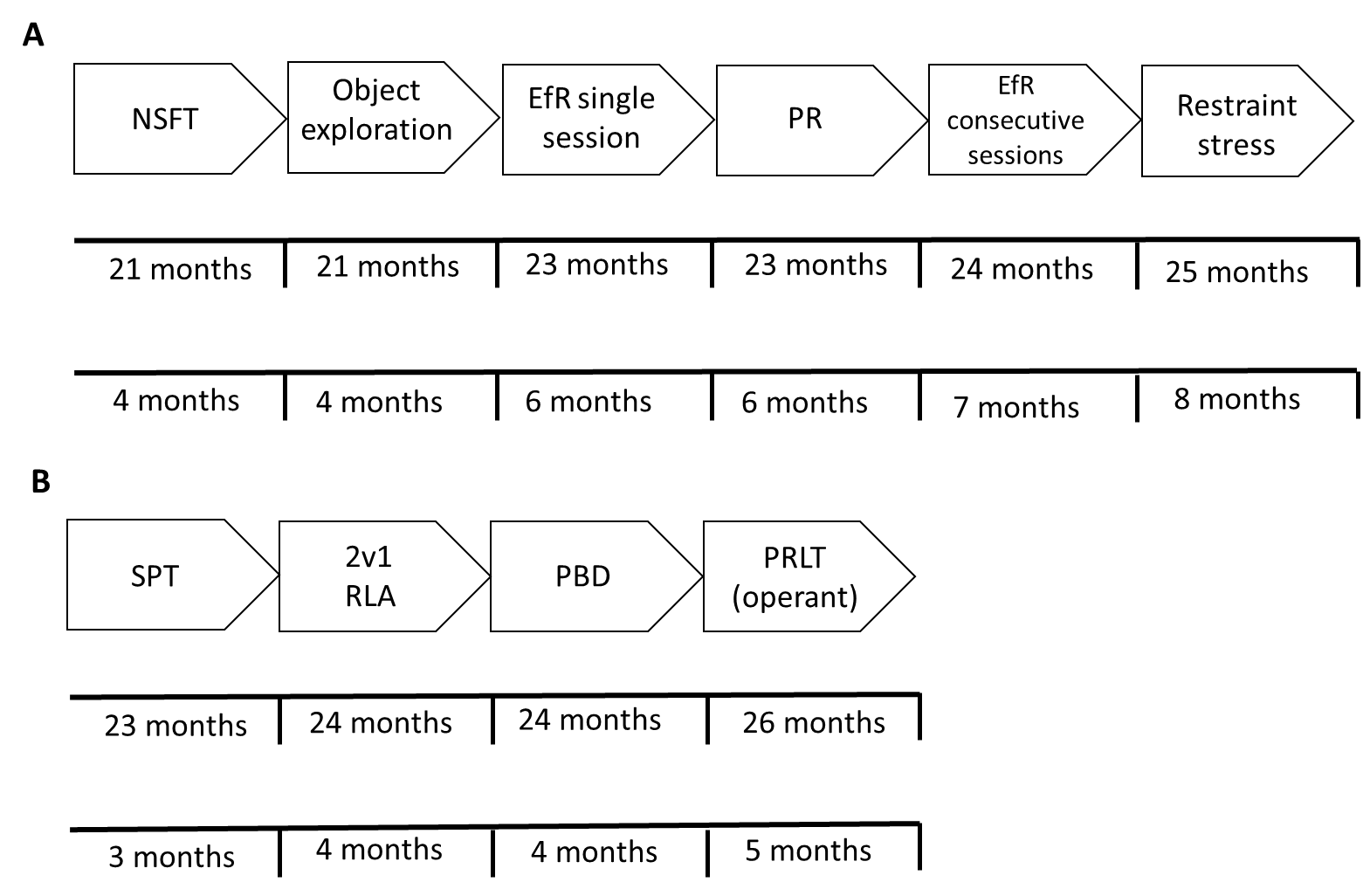

***S1. Timeline for analysis of age-related behaviours.*** *Two cohorts of rats underwent a series of behavioural tasks.* ***A*** *Cohort 1.* ***B*** *Cohort 2. NSFT-novelty supressed feeding test, EfR- effort for reward, PR-progressive ratio, SPT- sucrose preference test, RLA-reward learning assay, PBD-probabilistic bowl digging, PRLT-probabilistic reversal learning.*

**S2**

| Experiment | Measure | Statistical exclusion in dataset |
| --- | --- | --- |
| Reward learning assay | Reward bias | N = 1 young, n = 1 aged outliers. |
| Probabilistic bowl digging | Trials to first rule | None |
|  | Number of reversals | None |
|  | Win-stay probability | None |
|  | Lose shift-probability | N = 1 young, acquisition phase |
|  | % Sucrose preference | N = 1 aged |
| Progressive ratio | Food res- final FR completed | N = 1 young |
|  | Food res-breakpoint | None |
|  | Ad lib- breakpoint | None |
| Effort for reward | First test-trials | N = 1 young |
|  | First test-chow | None |
|  | Food res-trials | N = 1 young, n = 1 old full exclusion |
|  | Food res-chow | N = 1 aged (session 5) |
|  | Ad lib-trials | N= 1 young (session 5), n = 1 aged full exclusion |
|  | Ad lib-chow | N = 1 aged (session 2) |
| Probabilistic reversal learning (operant) | Trial first rule was learned | N = 2 young (session 4, 7)  N = 1 aged (session 6) |
|  | Reversals | N = 1 young (session 8) |
|  | Win-stay | N = 3 young (Session 4, 5, 6), n = 1 aged (session 3). |
|  | Lose-shift | N = 4 young (session 1, 2, 3, 7) and n = 1 aged, full exclusion |
|  | Initiation time | N = 2 young (session 3 and full exclusion) and n = 2 aged (session 1 and full exclusion). |
| Novelty supressed feeding test | Latency to approach | None |
|  | Latency to eat | None |
|  | Amount consumed | None |
| Restraint stress | Plasma CORT | N = 1 young (missing value) |
| Position based object exploration | Time spent exploring | N = 1 aged |
|  | Bouts of exploration | N = 1 young, n = 1 aged |
|  | % preference for side | N = 1 young |

***S2*** *Summary of excluded data points across experiments according to criteria outlined in methods.*
