## Supplementary methods for "Characterisation of behaviours relevant to apathy syndrome in the aged male rat"

**Digging training**

Habituation: On the first day, cage mates were placed in a 40 cm^2^ arena with a tray liner together covering the bottom and two pottery bowls placed at the back of the arena, approx. 1cm apart from each other. Each bowl was filled with reward pellets (LabDiet, 45 mg). Rats explored the arena with the lid fully closed for 10 mins. On the second day, rats were placed individually in the arena for 10 mins.

Pretraining: one reward pellet was placed in either the 10 o’clock, 12 o’clock or 2 o’clock position in one of the bowls. Once the rat had explored the bowls and found the pellet, a pellet was placed in the left-hand corner to encourage the rat to disengage from the bowl and return to the front of the arena. This was repeated until 12 trials were completed. Position of the baited bowl was presented in a pseudorandom order, and this applied to all stages of training. If the rat failed to find the pellet after 30 secs, the trial was scored as an omission. There were two sessions of pretraining.

Digging training: In the first trial, a pellet was placed in one bowl as described in the previous stage. In the second trial, the same bowl was filled with 1 cm sawdust and a pellet was buried in the same position. This was repeated for 12 trials. On the second and third day, this stage was repeated but with 2 cm sawdust and the empty bowl was removed after the rat started digging in the baited bowl.

Discrimination training: The rat was presented with two bowls filled with 2 cm of a different digging substrate e.g., sponge, perlite, cardboard squares etc. One of these substrates was consistently baited with one reward pellet and the other was mixed with a crushed reward pellet to prevent odour discrimination. In the first trial, the rat explored both bowls until the pellet was found. In the second trial, as soon as the rat chose one bowl, the other was removed. If the rat chose the baited bowl, it was recorded as correct and if the non-baited bowl was chosen it was recorded as incorrect. This was repeated until the rat chose the baited bowl over 6 consecutive trials within a maximum of 20 trials. After this, the rat was considered fully trained and ready for testing.

**Radioimmunoassay**

To denature corticosteroid binding globulin, plasma was diluted 1:50 with citrate buffer (25 mM tri-sodium citrate, 50 mM sodium dihydrogen orthophosphate and 1g/L bovine serum albumin (BSA) (Sigma-Aldrich, UK), pH 3.0). 100 ng/ml of CORT was serially diluted 1:2 in citrate buffer to create an 11-point standard curve. Known concentrations 20 ng/ml and 100 ng/ml CORT were used as quality controls (QCs). Rabbit anti-rat corticosterone primary antibody (a kind gift from G. Makara, Institute of Experimental Medicine, Budapest, Hungary) was diluted 1:50 in 0.2 ml distilled H2O, 9.8 ml 0.9 % saline solution, 500 ml citrate buffer and added to each sample, the QCs and the standard curve. Iodine corticosterone tracer ([^125^I], Izotop Institute of Isotopes Co., Ltd., Hungary) was then added, diluted to give between 3500-4000 counts per minute (cpm). Samples, QCs and standards were gently shaken and then stored overnight at 4 ͦC to incubate.

Following overnight incubation, samples, QCs and standards were precipitated with charcoal solution (0.05 g Dextran T70 (Pharmacia Biotech, Sweden) and 0.5 g charcoal (Fluka Analytical, Sigma, UK) per 100 ml citrate buffer). They were then centrifuged for 15 min at 4000 rpm at 4 ^ͦ^C. Supernatant was aspirated, and the remaining charcoal pellet was analysed using a gamma counter (E5010 Cobra II Auto Gamma, Perkin Elmer, Netherlands).
